# Efficient Genome Editing with Chimeric Oligonucleotide-Directed Editing

**DOI:** 10.1101/2024.07.09.602710

**Authors:** Long T. Nguyen, Noah R. Rakestraw, Brianna L.M. Pizzano, Cullen B. Young, Yujia Huang, Kate T. Beerensson, Anne Fang, Sydney G. Antal, Katerina V. Anamisis, Coleen M.D. Peggs, Jun Yan, Yangwode Jing, Rebecca D. Burdine, Britt Adamson, Jared E. Toettcher, Cameron Myhrvold, Piyush K. Jain

## Abstract

Prime editing has emerged as a precise and powerful genome editing tool, offering a favorable gene editing profile compared to other Cas9-based approaches. Here we report new nCas9-DNA polymerase fusion proteins to create chimeric oligonucleotide-directed editing (CODE) systems for search-and-replace genome editing. Through successive rounds of engineering, we developed CODEMax and CODEMax(exo+) editors that achieve efficient genome modifications in human cells with low unintended edits. CODEMax and CODEMax(exo+) contain an engineered Bst DNA polymerase derivative known for its robust strand displacement ability. Additionally, CODEMax(exo+) features a 5’ to 3’ exonuclease activity that promotes effective strand invasion and repair outcomes favoring the incorporation of the desired edit. We demonstrate CODEs can perform small insertions, deletions, and substitutions with improved efficiency compared to PEMax at many loci. Overall, CODEs complement existing prime editors to expand the toolbox for genome manipulations without double-stranded breaks.

## Introduction

Prime editing has gained prominence as a highly effective genome editing tool due to its precision and versatility^1-3^. Pioneered by Anzalone *et al*.^1^, the technology can correct all 12 possible single-base substitutions and make small insertions or deletions without generating double-stranded breaks. Prime editing typically uses a nickase Cas9-reverse transcriptase (nCas9-RT) fusion protein (prime editor) and a prime editing guide RNA (pegRNA), which has a 3’ extension containing a primer binding site (PBS) and reverse transcription template (RTT) encoding the desired edit. The 5’ spacer region of the pegRNA directs the prime editor to the target site, resulting in nicking of the non-target strand by nCas9. Hybridization of the 3’ PBS to the nicked strand then generates an initiation site for reverse transcription to occur. Subsequently, the correct edits are installed into a 3’ edited DNA flap which can be incorporated into the genome^1,4^.

Although prime editing has been iteratively improved through many rounds at the level of both the prime editor and pegRNA, the system still suffers from variable efficiency across genomic loci^5-7^. Existing prime editors use M-MLV or other reverse transcriptases, which possess relatively moderate processivity, hindering their capability to perform more complex edits effectively. Additionally, after the prime editor generates a 3’ edited flap, it competes with the 5’ unedited flap for incorporation into the genome, reducing the likelihood of a successful edit. Furthermore, since the pegRNA contains a PBS that is complementary to the spacer sequence, it forms a stable RNA-RNA duplex, which creates a barrier for ribonucleoprotein complexation between the pegRNA and the prime editor^8,9^. Recent efforts have been made to circumvent the auto-inhibitory interaction within the pegRNA by optimizing the melting temperature of the PBS and by incorporating mismatches into the PBS to reduce misfolded pegRNA interactions which leads to improved prime editing efficiency^8,9^. Although prime editing can make versatile edits without induction of double-strand breaks in the DNA, the accuracy of prime editors still has room for improvement. One of the most common imprecise edits observed for prime editing is an overextension of the reverse transcriptase past the RTT into the scaffold region of the pegRNA. This readthrough becomes problematic because it results in the incorporation of undesired bases into the genome^10^, although recent studies have shown methods to lower its occurrence by engineering highly structured regions within the pegRNA^4,11^.

We reasoned that creating chimeric oligonucleotide-directed editing (CODE) systems consisting of a DNA-dependent DNA polymerase paired with a chimeric pegRNA (cpegRNA) containing a DNA primer binding site and a DNA polymerase template may address some of the limitations of current reverse transcriptase-based prime editors. We hypothesized that a cpegRNA could reduce the auto-inhibitory effect observed for traditional pegRNAs, as the DNA-RNA duplex is inherently less stable than the RNA-RNA duplex^12^. Moreover, DNA polymerases are an abundant family of proteins with diverse molecular properties, rendering them intriguing candidates to achieve chimeric oligonucleotide-directed editing. The use of DNA polymerases with advantageous properties, such as high processivity, proofreading capability, and minimal reverse transcriptase activity has the potential to improve editing efficiency, reduce unintended edits, and enable new types of edits to be performed.

Recent studies have also investigated the use of DNA-dependent DNA polymerases in place of reverse transcriptases for precise search-and-replace genome editing^13,14^. Liu *et al*. reported a four-component DNA Polymerase Editing system comprised of nCas9, Phi29 DNA polymerase-MCP fusion protein, sgRNA, and a chimeric DNA-RNA template with an MS2 loop. Ferreira *et al*. developed Click Editors that utilize nCas9 fused to both a DNA polymerase and a ssDNA tethering domain that localizes the DNA template for polymerization to the edit site. These studies demonstrate the potential of utilizing DNA-dependent DNA polymerases as a new tool for precision genome editing.

Here we survey a new class of 13 CODEs that consist of a cpegRNA and a nickase Cas9-DNA polymerase fusion protein. This simple two-component system allows for delivery via plasmids or ribonucleoprotein (RNP) complexes for effective and accurate genome editing. We observed that CODE improved gene correction efficiency compared to conventional PE2 and PEMax at several genomic loci with low unintended scaffold incorporation. Overall, engineered CODEs expand the gene editing toolbox and offer versatility as well as flexibility toward therapeutic applications.

## Results

### A survey of 13 DNA polymerase-mediated editors for targeted genome modification

To determine whether a nickase Cas9 (nCas9) fused to a DNA polymerase (DNAP) could perform precise genome editing when delivered as a ribonucleoprotein complex, we first constructed nCas9 fusion proteins paired with a variety of wild-type polymerases from viral or bacterial origins (Fig. 1a). A total of 13 DNAPs with diverse properties such as thermostability, proofreading activity, processivity, and size were selected and screened (Fig. 1b). Each construct was expressed in *E. coli* and purified for ribonucleoprotein delivery. To evaluate the efficiency of CODE candidates, we generated a HEK293T-based reporter cell line containing an open reading frame with a green fluorescence protein (GFP) upstream of a red fluorescence protein (mCherry). We also installed a premature termination codon (PTC) into the mCherry gene so that the cells only display green fluorescence, similar to Anzalone *et al*^15^. Upon successful A-to-G substitution within the PTC, the stop codon is converted to the CAA codon which allows for mCherry translational readthrough.

**Figure 1.**
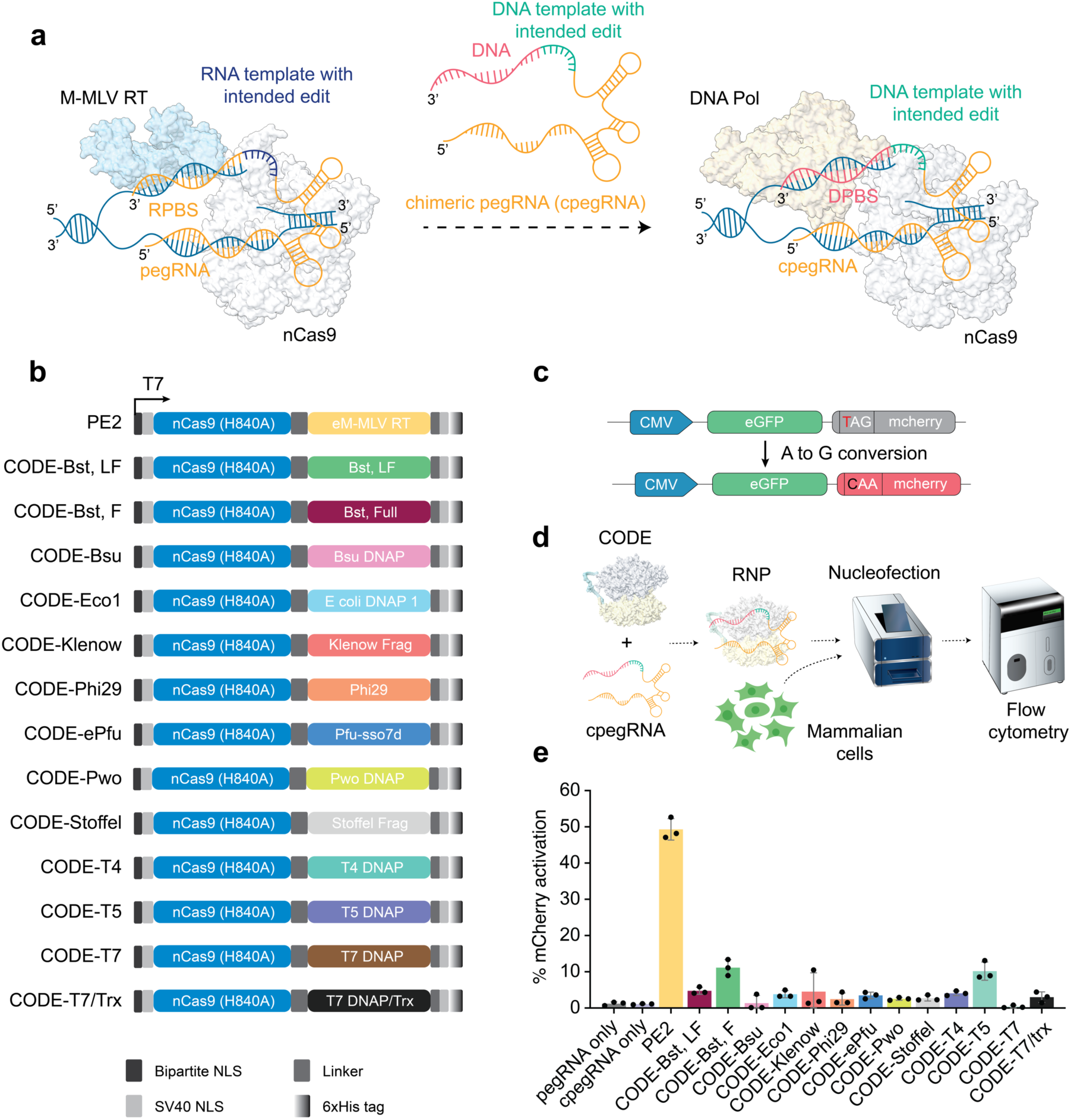
Initial screening for activity of CODE candidates. (**a**) Schematic of the development of CODEs. CODEs consist of a nCas9-DNAP fusion protein and a chimeric pegRNA (cpeg) containing a guide RNA and ssDNA template with intended edits and primer binding site (PBS). (**b**) Architecture of bacterial expression plasmids of CODEs. The editor expression is driven by T7 promoter, and 6x Histidine tag is located at the C-terminus is employed for purification purposes. (**c**) Construction of HEK293T reporter cell line supporting base conversion via prime editing or CODE. (**d**) Schematic of the workflow for nucleofection of CODEs and cpegRNA into HEK293T reporter cell line. (**e**) Percentage of mCherry activation of CODE candidates and the control engineered PE2 system. Error bars represent ± SD, where n = 3 biological replicates.

Next, we designed a cpegRNA to target the constitutively expressed GFP-mCherry reporter gene containing the PTC. The cpegRNA is comprised of a 20-nt RNA targeting sequence, a guide RNA scaffold, and a 3’-end DNA extension sequence containing a primer binding site (PBS) and a DNAP template encoding the desired corrections (Fig. 1c). Each CODE candidate was complexed with the cpegRNA prior to delivery into HEK293T cells via nucleofection and edit efficiency was quantified by the percentage of mCherry positive cells (Fig. 1d)^10^.

We observed that many CODE candidates were able to induce mCherry activation, albeit with low efficiencies compared to the engineered prime editor PE2. Among the most efficient wild-type CODEs were those that employed polymerases from T4 and T5 bacteriophages, reaching 4.1% and 10.2% mCherry activation, respectively. Additionally, the large fragment Bst DNAP derived from *Geobacillus stearothermophilus* achieved 4.7% while full length Bst DNAP was even more effective at 11.1% mCherry activation. Liu *et al*.^13^ and Ferreira da Silva *et al*.^14^ recently reported prime editing adjacent systems that utilize DNA-dependent DNA polymerases such as Phi29 DNAP and Klenow fragment. Within our editing system, we observed that CODE candidates employing DNA Polymerase I and the Klenow fragment exhibited moderate mCherry activation (3.8% and 4.5%, respectively) while Phi29 DNAP showed low editing activity (2.3%) (Fig. 1e). Although our chimeric editing system differs from these studies, our data support the reported abilities of Phi29 and the Klenow fragment to extend primed templates at a nick in the genomic DNA. Together, these data demonstrate that multiple DNA polymerases can utilize a DNA template within a chimeric pegRNA to perform precise edits in mammalian cells.

### Engineering T4 DNA polymerase for improved editing efficiency

For further engineering of CODEs for greater functionality and utility, we selected T4 DNAP and Bst, large fragment DNAP (Bst-LF). T4 DNAP is a mesophilic polymerase that possesses strong 3’-5’ exonuclease activity^16,17^. We strived to determine how the T4 DNAP location impacted the CODE-T4 editing efficiency by positioning the T4 DNAP either on the C-terminus (CODE-T4v1) or the N-terminus (CODE-T4v2) of nCas9 connected by a 33-amino acid linker (Fig. 2a). We did not observe much difference in the percentage of mCherry activation between the N-terminal and C-terminal fusion constructs. We next aimed to investigate how the 3’-5’ exonuclease activity of T4 DNAP affected the performance of CODE (Fig. 2b). We installed a Y320A mutation on T4 DNAP of CODE-T4v1 editor (CODE-T4v3), which has been shown to diminish the polymerase exonuclease activity by 50-fold^18^. Notably, this single mutation increased the efficiency of mCherry activation 2.4-fold compared to the wild-type editor (Fig. 2b). We proceeded to install a T4 Gene 32 Protein (gp32) on the N-terminus of T4 DNA polymerase to generate a CODE-T4v4 editor (Fig. 2a,b). T4 gp32 is a single-stranded binding protein (SSB) that is crucial for T4 replication and repair^19,20^. However, we observed reduced activity compared to CODE-T4v3, possibly due to gp32 SSB being inactive in a fusion format or being sterically hindered by nCas9 and T4 DNAP.

**Figure 2.**
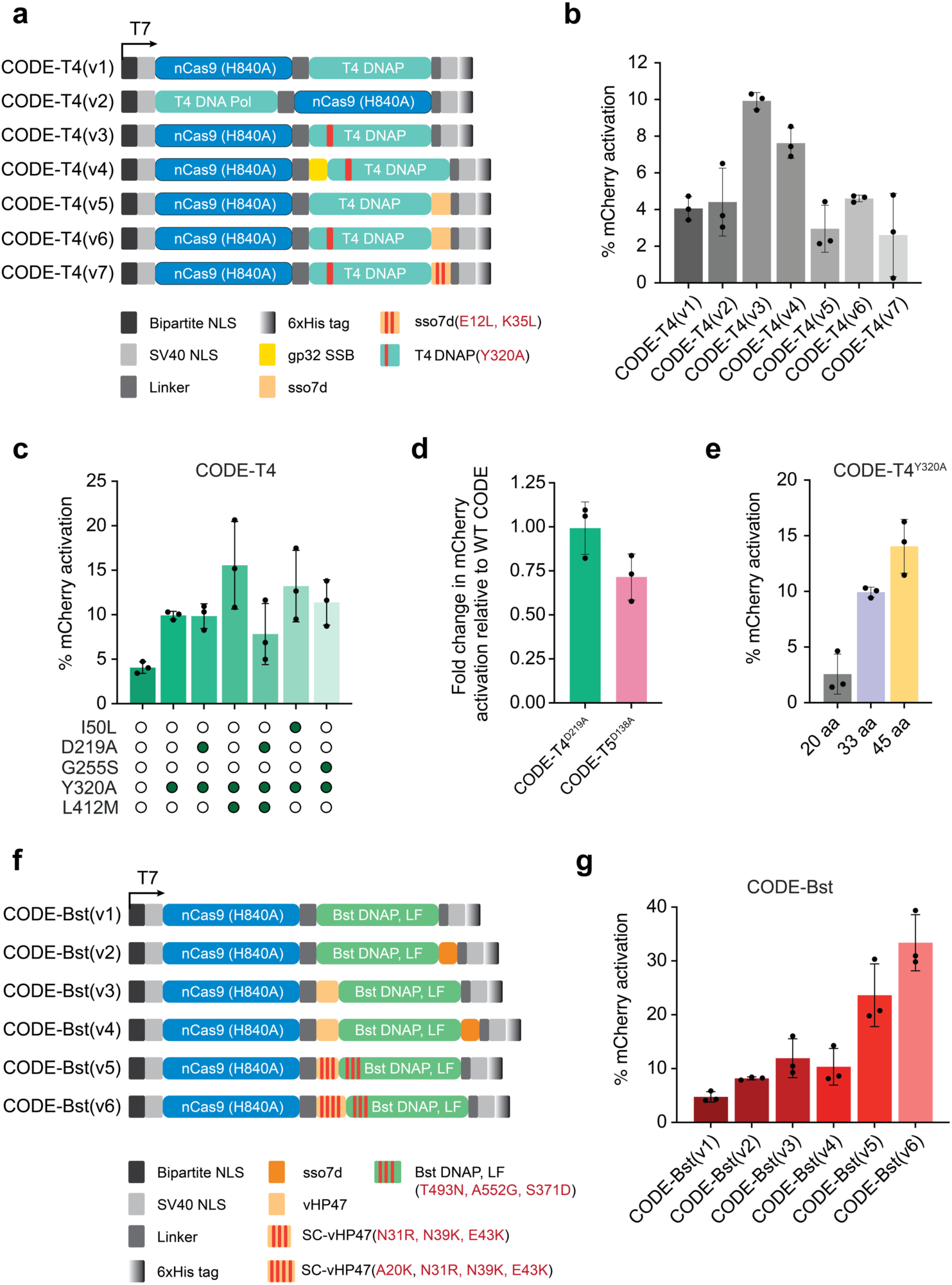
Engineering of T4 and Bst chimeric oligonucleotide-directed editors for improved editing. (**a**) Architecture of engineered CODE-T4 editors with domain rearrangement strategies. (**b**) Percentage of mCherry activation of the CODE-T4 variants in (**a**). (**c**) Engineering attempts to alter the T4 DNAP processivity and fidelity to create improved CODE-T4 variants beneficial mutations. (**d**). (**e**) Optimization of amino acid linker length between nCas9 and T4 DNAP in the fusion construct. (**f**) Engineering attempts to alter the Bst-LF DNAP thermostability to create improved CODE-Bst variants with beneficial mutations. (**g**) Percentage of mCherry activation of the CODE-T4 variants in (**f**). Error bars represent ± SD, where n = 3 biological replicates.

We then inserted a sso7d DNA binding domain at the C-terminus T4 DNAP of CODE-T4v1 and CODE-T4v3 to create CODE-T4v5 and CODE-T4v6. Sso7d, a DNA binding protein derived from *Sulfolobus solfactaricus*, is known to greatly enhance the processivity of DNA polymerases^21^. We sought to understand if this binding domain could improve our CODE-T4 editors. Interestingly, we did not observe any significant mCherry activation of these two CODEs compared to the original CODE-T4v1. The CODE-T4v6 editor, which possesses a Y320A mutation, exhibited a similar editing efficiency to the CODE-T4v1. Since the sso7d DNA binding domain possesses some ribonuclease activity, we reasoned that it could potentially degrade the cpegRNA and have a negative impact on CODE^22^. We therefore deactivated this ribonuclease activity by introducing two mutations E12L and K35L into the sso7d domain to generate CODE-T4v7. However, this version of CODE performed poorly compared to CODE-T4v3 with a 3.8-fold decrease in efficiency (Fig. 2b).

We next focused on engineering the T4 DNAP itself, which has been well-studied and extensively characterized. Mutations such as L412M and I50L have been shown to increase the processivity, although these substitutions cause a slight increase in replication errors^23,24^. We therefore generated combinations of these CODE-T4 mutants and tested them in HEK293T cells. Notably, we observed up to 20.7% mCherry activation for the CODE-T4^Y320A/L412M^ mutants, which is a 3.8-fold increase compared to CODE-T4v1. We also noted a boost in efficiency for the CODE-T4^I50L/Y320A^ and CODE-T4^G255S/Y320A^ mutants compared to the original CODE-T4v1 (Fig. 2c).

We hypothesized that the 3’-5’ exonuclease activity of T4 DNA polymerase might have a negative impact on the editing efficiency as it could remove bases at the newly synthesized 3’-flap^25,26^. To test this hypothesis, we generated a CODE-T4^D219A^ and CODE-T5^D138A^ with deficient 3’-5’ exonuclease activity and compared them against the corresponding wild-type CODE-T4 and CODE-T5. Interestingly, we observed no difference in mCherry activation in HEK293T reporter cells, indicating that this 3’-5’ exonuclease activity might have minimal involvement in the prime editing process (Fig. 2d). We finally optimized the amino acid linker length between the nCas9 and T4 DNAP and found that a 45 amino acid linker exhibited the best mCherry activation efficiency (Fig. 2e).

### Engineering Bst DNA polymerase for improved editing efficiency

Bst-LF DNAP is a thermophilic DNA polymerase that has strong strand displacement activity; therefore, Bst-LF is often used in isothermal amplification technologies such as Loop-mediated isothermal amplification (LAMP)^27,28^. Although wild-type Bst-LF is optimally active at high temperatures in amplification reactions, we observed moderate editing efficiency by Bst-LF and full length Bst DNAP in HEK293T cells. Additionally, prime editor 2 (PE2), which utilizes an engineered M-MLV variant containing five-point mutations that increase the thermostability and processivity, was shown to achieve dramatic increase in prime editing efficiency compared to prime editor 1 (PE1) that contains a wild-type M-MMLV reverse transcriptase^1^. While Bst-LF DNAP is naturally a thermophilic enzyme, we reasoned that enhancing its thermostability further might improve its overall performance inside cells. We therefore explored multiple approaches to engineer Bst-LF mutants within our CODE systems to see if increasing the thermostability of Bst-LF DNAP would increase its activity in cells.

Paik *et al*.^*29*^, used a machine learning approach to generate a Bst DNAP variant with increased thermostability. The Bst-LF DNAP variant, referred to as Br512, consists of a modified 47 amino acid actin-binding protein called villin headpiece fused to the N-terminus of the Bst-LF. The fusion of the villin headpiece to Bst-LF DNAP is hypothesized to improve protein folding and increase processivity via stabilization of the DNA/protein complex. We employed the Br512 DNAP in CODE (referred to as CODE-Bstv3) and observed a nearly 3-fold increase in mCherry-positive cells compared to wild-type Bst-LF. In an attempt to further increase efficiency, we tested additional variants of Br512 engineered to have even stronger thermostability. Br512g3.1 and Br512g3.2 variants differ from Br512 in the villin headpiece, where point mutations were rationally designed to supercharge and stabilize the domain (referred to as SC-vHP47). The Br512g3.1 variant bears 3 mutations (N31R, N39K, E43K) on the SC-vHP47, whereas the Br512g3.2 bears 4 mutations (A20K, N31R, N39K, E43K). Furthermore, these two Bst-LF DNAP variants bear additional mutations within the polymerase domain (T493N, A552G, S371D), resulting in a significant increase in thermostability over Br512 while maintaining high functionality in LAMP reactions up to 74°C^29^. By incorporating these Br512g3.1 and Br512g3.2 DNAP variants into our chimeric oligonucleotide-directed editors, we generated the CODE-Bstv5 and CODE-Bstv6 systems. Notably, CODE-Bstv5 and CODE-Bstv6 resulted in 23.6% and 33.4% efficiency installing the desired edits, approximately 5-fold and 7-fold increases in mCherry activation compared to the original CODE-Bstv1 system.

### In-house synthesis of chimeric pegRNA improves editing efficiency

Traditionally, a pegRNA consists of 100% RNA bases and therefore can be synthesized either by enzymatic or chemical synthesis reactions. This advantage provides flexibility to deliver prime editing systems into cells as well as animal models. Oftentimes, it is convenient to co-deliver plasmids encoding prime editors and pegRNAs driven by a U6 promoter. Since the cpegRNA is a chimeric entity consisting of both RNA and DNA bases, it cannot be synthesized by naturally occurring enzymes. Instead, cpegRNAs must be chemically synthesized, which poses challenges for delivery approaches and associated synthesis costs.

To address this hurdle, we developed an inexpensive ligation-based method to synthesize cpegRNAs. Taking advantage of T4 RNA ligase I, which can ligate a 3’-hydroxyl RNA to a 5’-phosphorylated DNA, we established a synthesis protocol for generating full length cpegRNA (Fig. 3a). A benefit of this ligation reaction is that one can easily make a multitude of edit types at the same genomic locus by modifying single stranded DNA oligos which each utilize the same single-guide RNA. This approach drastically decreases the synthesis cost compared to synthetic cpegRNA. Additionally, we placed four modified 2′-O-methylated uracils at the end of the single guide RNA to improve stability and reduce sgRNA scaffold incorporation in the cells (Fig. 3b). We tested in-house synthesized cpegRNAs with CODE-Bstv6 and noted an improved performance with 43.7% mCherry activation (Fig. 3c,d).

**Figure 3.**
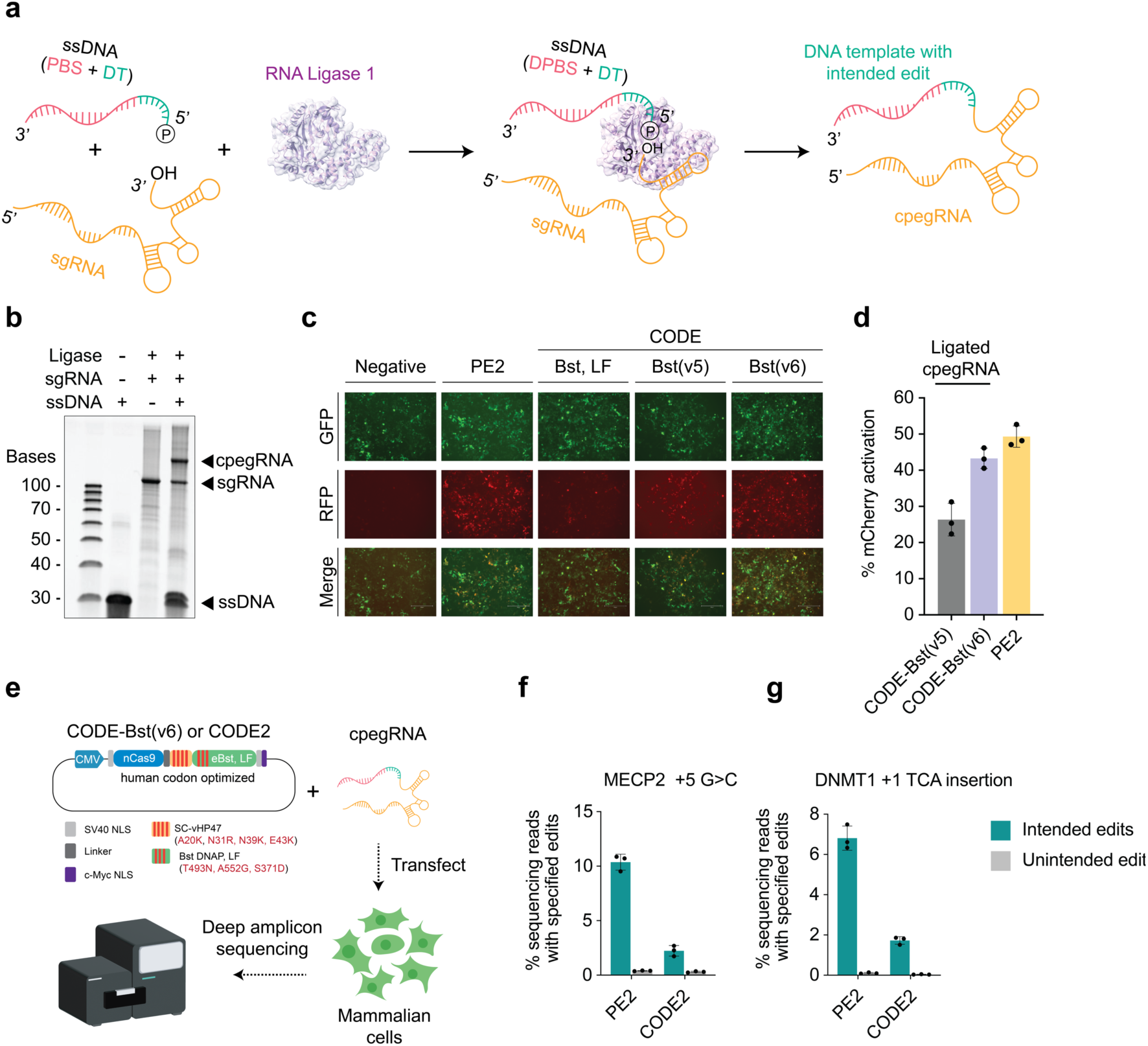
In-house synthesis of cpegRNA ligation reaction. (**a**) Schematic of the T4 RNA Ligase I-mediated cpegRNA synthesis. (**b**) Representative of denaturing gel showing successful ligation of sgRNA and ssDNA oligo to generate cpegRNA that targets mCherry gene. (**c**) Visualization of HEK293T cells by fluorescence microscopy showing the mCherry activation by PE2 and engineered CODE-Bst variants with ligated cpegRNA. Cells were transfected with prime editors and CODEs 72 hours prior to imaging. (**d**) Quantification of mCherry activation in (c) via flow cytometry. (**e**) Schematic of the workflow for transfection of plasmid encoding human codon-optimized CODE and synthetic or ligated cpegRNA. (**f**) and (**g**) Efficiency of intended and unintended modifications of PE2 and CODE2 at *MECP2* and *DNMT1* loci, respectively. Error bars represent ± SD, where n = 3 biological replicates.

### Efficient editing of endogenous gene loci with CODEMax and CODEMax(exo+)

We next aimed to investigate how efficiently CODE performs when targeting endogenous genes. Using nucleofection strategies, we observed a significant reduction in editing efficiency for our CODE-Bstv6, possibly due to the cpegRNA chimeric nature initiating cellular response, leading to its degradation. We then explored alternative methods to deliver CODEs by comparing RNP, mRNA, and plasmid approaches. We proposed that encapsulating the cpegRNA with transfection reagents could provide it with some protection from the cellular environment. After several rounds of optimization, we showed that plasmids encoding CODEs and synthetic cpegRNAs can be co-delivered into mammalian cells via transfection (Fig. 3e-g and Fig. S1).

To enhance editing efficiency, we adopted two additional nCas9 mutations (R221K and N394K) that were developed by Chen *et al*^30^. to convert PE2 into PEMax. We called this version of chimeric oligonucleotide-directed editing CODEMax, which is a combination of the engineered CODE-Bstv6 and a mutated nCas9 variant (R221K and N394K). Furthermore, our initial screening of CODE candidates (Fig. 1e) showed better performance of a chimeric oligonucleotide-directed editor using full-length Bst DNAP compared to that with Bst-LF. Interestingly, the difference between the truncated and full-length polymerase is the absence of a 5’-3’ exonuclease domain. We hypothesized that a polymerase that possesses strong 5’-3’ exonuclease activity may improve the editing efficiency (Fig. 4a,b and Fig. S2). In prime editing-like systems, the 3’ flap generated after extension by the polymerase, enters competition for incorporation into the genome with the 5’ unedited flap^31^. Therefore, having a polymerase with 5’-3’ exonuclease activity that can displace and degrade the 5’ unedited strand during extension may be beneficial (Fig. 4b). In support of this hypothesis, Liang *et al*.^32^ has demonstrated that the fusion of a T5 exonuclease at the N-terminus of the M-MLV reverse transcriptase increased efficiency of the PE2 system in plants. Therefore, we incorporated the 5’-3’ exonuclease domain back into CODEMax, hereafter referred to as CODEMax(exo+).

**Figure 4.**
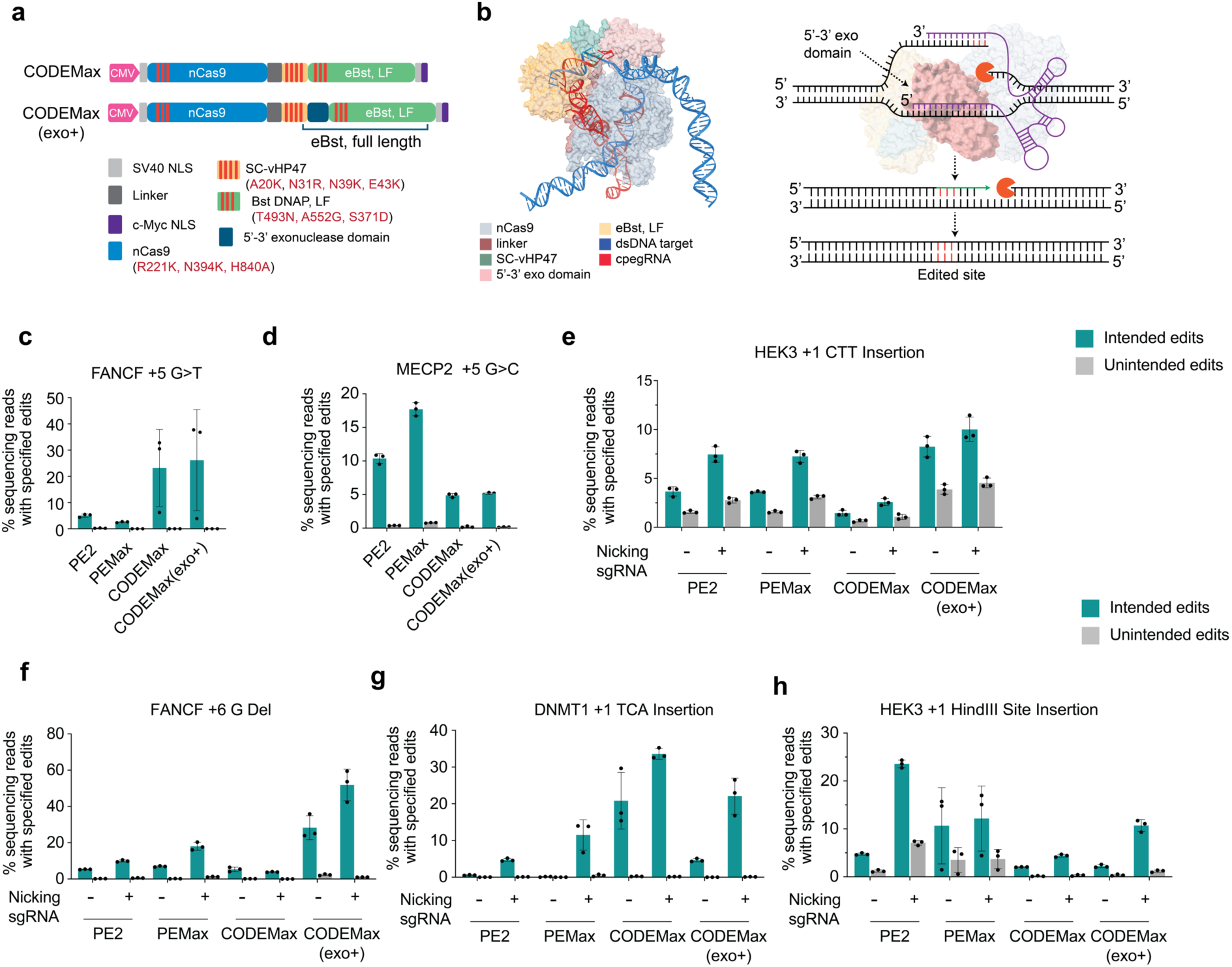
Efficient chimeric oligonucleotide-directed editing of endogenous gene loci with CODEMax and CODEMax(exo+). (**a**) Architecture of plasmid encoding CODEMax and CODEMax(exo+). (**b**) Alphafold3 predicted structure of the CODEMax(exo+) in complex with cpegRNA and target dsDNA^36^. (**c**)-(**h**) Endogenous targeting of CODEMax and CODEMax(exo+) at various gene loci in comparison with PE2 and PEMax. Error bars represent ± SD, where n = 3 technical replicates.

We tested CODEMax and CODEMax(exo+) by targeting multiple genomic regions with a variety of edit types such as base conversion and transversion, short insertion, and short deletion. We observed enhanced editing efficiency of CODEMax and CODEMax(exo+) compared to PE2 and PEMax at several loci such as *EMX1, FANCF, SRD5A3* and *DNMT1* with HEK3 and *MECP2* being exceptions. (Fig. 4c-h and Fig. S3-S5). It should be noted that these data were head-to-head comparisons of synthetic pegRNA/cpegRNA and protein-encoding plasmid via transfection. However, we noticed that expression of pegRNA under U6 promoter resulted in higher efficiency for PE2 and PEMax compared to delivery of synthetic pegRNA (Fig. S6). With the addition of a nicking guide, which significantly increases edit efficiency for traditional prime editors by evading mismatch repair (MMR), we also observed improved prime editing for CODE systems, suggesting a conserved presence of heteroduplex intermediates^1^ (Fig. 4e-h and Fig. S4). Additionally, CODEMax(exo+) outperformed CODEMax at a majority of edit sites. Lastly, we demonstrate that CODEMax and CODEMax(exo+) exhibited low unintended scaffold incorporation like that seen for PEMax at certain edit sites (Fig. S7). On the other hand, comparable amounts of total unintended edits were observed between PE and CODE systems (Fig. S8).

## Discussion

Together with two recent studies by Liu *et al*.^*13*^ and Ferreira da Silva *et al*.^14^, this study demonstrates that chimeric oligonucleotide-directed editors are effective genome editors but with a distinctly new approach in protein engineering and cpegRNA design, establishing a new class of Cas9-based editing tools. Our two-component CODE system was developed by screening 13 diverse polymerases and selecting the best candidates for further engineering, resulting in a thermophilic Bst DNA polymerase with a robust strand displacement capability and 5’-3’ exonuclease activity. We hypothesized that the strand displacement property promotes genome strand invasion during R-loop formation of nickase Cas9 at the target site, allowing for enhanced polymerization. Additionally, the 5’-3’ exonuclease activity of the Bst DNAP supports the degradation of the 5’-flap which leverages the incorporation of the newly polymerase-mediated extension of the 3’-flap into the genome, favoring the edit-incorporating outcomes.

The engineered CODE systems allow for further development of a whole new class of prime editing systems utilizing DNA-dependent DNA polymerases. DNA polymerases are abundant and diverse across all three domains of life. The specific properties of wild-type DNA polymerases that can be beneficial for prime editing like systems include thermostability, processivity, proofreading ability, and 5’ to 3’ exonuclease activity. As we have shown, further engineering of wild-type DNA polymerases can improve editing outcomes, but engineering efforts may also be directed towards specific applications. For example, a highly processive DNA polymerase may be required for longer insertions; however, for simpler edits, a polymerase with higher fidelity may be favored. Doman *et al*. have shown that different reverse transcriptase proteins perform better depending on the edit type and location^11^. Having a diverse toolbox of prime editors utilizing reverse transcriptase or DNA polymerase-based editors enables broad applications for genome engineering.

## Limitations of the Study

Although we tested the CODEs across multiple genomic loci, further engineering and validation are needed to explore what additional benefits the CODE systems can offer. Many of the limitations possessed by traditional reverse transcriptase-based editors are not completely addressed by CODE systems. The major limitation still encountered by CODEs is diverse editing outcomes depending on the loci and edit sites. Although many wild-type DNA polymerases were screened and subjected to engineering, additional engineering can be performed to continue to improve effectiveness. Moreover, CODEs still are unable to effectively perform long insertions and deletions. However, different polymerases or the adoption of twin editing or integrase-mediated gene insertion strategies may benefit from using CODEs^33,34^. Lastly, compared to pegRNA, cpegRNA are more expensive to synthesize and less versatile. For this reason, further development of effective delivery methods of cpegRNA is worthwhile.

## Materials and Methods

### General cloning methods and plasmid construction

CODE gene fragments were either obtained from Addgene or synthesized by Twist Biosciences. Bacterial and mammalian expression plasmids for CODEs were cloned using In-Fusion® cloning (Takara Bio, Cat# 638948). Q5 high-fidelity polymerase (New England Biolabs, Cat# M0491L) was used to amplify gene fragments and non-lentiviral backbone for cloning as well as genomic DNA for deep sequencing. For In-Fusion® cloning involving the assembly of lentiviral backbones, PrimeSTAR® GXL DNA Polymerase (Takara Bio, Cat# R050A) was used for amplification. For transfection into mammalian cells, plasmids expressing CODEs were prepared using ZymoPure II Plasmid Midiprep kit (Zymo Research, Cat# D4201) and diluted down to 1 mg/μL prior to transfection.

### Chimeric pegRNA synthesis via T4 RNA ligase-mediated ligation

All pegRNAs, sgRNAs, and 5’ phosphorylated ssDNAs were purchased from IDT. cpegRNAs were also purchased from IDT unless otherwise indicated that they were produced via ligation of 5’ phosphorylated ssDNA to sgRNAs. Ligations were prepared using T4 RNA Ligase I (New England Biolabs, Cat# M0204) as follows: 3.5 μL T4 RNA Ligase Buffer, 4.5 μL PEG8000, 2 μL T4 RNA Ligase I, 3 μL 10mM ATP, 2 μL 100uM sgRNA, 5 μL 100 mM 5’ phosphorylated ssDNA, 0.25 μL Murine RNase Inhibitor (New England Biolabs, Cat# M0314), and 14.75 μL water. Reaction volumes greater than 35μL led to decreased ligation efficiencies. Reactions were incubated at 16°C for 16 hours and purified with Monarch® RNA Cleanup Kits (New England Biolabs, Cat# T2030) according to manufacturer instructions. Products were analyzed via 10% TBE-Urea PAGE electrophoresis (Biorad, Cat# 3450088).

### CODE protein expression and purification

Bacterial expression plasmids carrying CODEs were transformed into homemade competent cells propagated from Rosetta™ 2(DE3)pLysS Singles™ Competent Cells (Millipore Sigma, Cat# 71401). Individual colonies were picked and inoculated in 50 mL Luria Broth (Fisher Scientific, Cat# BP9723-2) overnight at 37°C. The culture was then scaled up to 4-12 liters of Terrific Broth (RPI, T15000-10000.0) and grown until OD = 0.8-1.0. The culture was then quickly cooled on ice for 10-15 minutes and induced with 1 mM isopropyl ß-D-1-thiogalactopyranoside (IPTG) (Gold Biotechnology, Cat# I2481C100). For CODE constructs that were built based on the pET-PE2-His backbone (Addgene, #170103), the culture was induced at 18°C for 5 hours followed by 26°C for 14-18 hours. For CODE constructs that were built based on PE-Max-pET21a backbone (Addgene, #204471), the culture was induced at 18°C for 16-18 hours.

Cell pellets were collected the next day by centrifugation (4000 xg for 10 minutes), suspended in 100-150 mL lysis buffer (500 mM NaCl, 50 mM Tris-HCl pH = 7.5, 1 mM TCEP-HCl, 20 mM imidazole, and 5% glycerol) followed by sonication. The lysate was centrifuged at 40,000 xg for 45 minutes before passing through a 0.45 μm filter. The clarified lysate was then injected into a prepacked Ni-NTA affinity column (EconoFit Nuvia IMAC Column, Biorad #12009287) in a FPLC (NGC Quest Plus, Biorad) pre-equilibrated with lysis buffer. Proteins were eluted from the column with 40 mL of elution buffer (500 mM NaCl, 50 mM Tris-HCl pH = 7.5, 1 mM TCEP-HCl, 300 mM Imidazole, and 5% glycerol). The eluted solution was concentrated in an Amicon® Ultra Centrifugal Filter, 50 kDa MWCO (Millipore Sigma, UFC905024) down to 10-15 mL and equilibrated with 40 mL of Buffer A (200 mM NaCl, 50 mM Tris-HCl pH = 7.5, 1 mM TCEP-HCl, and 5% glycerol). The protein mixture was then passed through a 5 mL Hitrap Heparin HP column (Cytiva, Cat# 17040701) pre-equilibrated with Buffer A. The column then underwent gradient elution from Buffer A to Buffer B (2000 mM NaCl, 50 mM Tris-HCl pH = 7.5, 1 mM TCEP-HCl, and 5% glycerol). The purest fractions of the protein were pooled together and concentrated in an Amicon® Ultra Centrifugal Filter (50 kDa MWCO) in final buffer C (500 mM NaCl, 50 mM Tris-HCl pH = 7.5, 1 mM TCEP-HCl, and 5% glycerol) before storing at −80°C. When in use, the protein was diluted in storage buffer (300 mM NaCl, 10 mM Tris-HCl, 0.1 mM EDTA, 1 mM DTT, 50% glycerol, 0.1% Triton® X-100, final Ph = 7.4 at 25°C) down to 50 μM and store at −20°C.

### Mammalian cell culture

HEK293T and Lenti-X™ 293T were obtained from ATCC (CRL-3216) and Takara Bio (#632180), respectively. U2OS cells were obtained ATCC (HTB-96). The cells were tested with mycoplasma using MycoAlert® Mycoplasma Detection Kit (Lonza, Cat# LT07-118). The cells were cultured and passaged in D10 medium containing DMEM high glucose with GlutaMAX™ supplement and pyruvate (Gibco, Cat# 10569044), 10% Fetal Bovine Serum (Gibco, Cat# A3160902), 1X Penicillin-Streptomycin (Gibco, Cat# 15140122), and 1X MEM non-essential amino acids (Gibco, 11140035). All cell lines were incubated at 37°C and 5% CO^2^.

### Reporter cell line and stably expressed CODE cell line generation

For lentiviral packaging in a T75 flask, 10 μg of transfer plasmid was co-transfected with 5 μg of pMD2.G (Addgene, #12259) and 7.5 μg of psPAX2 (Addgene, #12260) into Lenti-X™ 293T cells with 50 μL of Lipofectamine 3000 and 42 μL P3000 Enhancer reagents (ThermoFisher, Cat# L3000008), which were diluted in Opti-MEM™ I Reduced Serum Medium (Gibco, 31985062) following the manufacturer instructions. The medium was changed to D10 medium six hours later, and the cells were incubated for an additional 48-60 hours. The cells were harvested and pelleted down via centrifugation at 4000 xg for 10 minutes at 4°C. The supernatant was passed through a 0.45 μm filter, aliquoted, and stored at −80°C until use.

For lentiviral transduction, 5 x10^5^ HEK293T cells were infected with multiple dilutions of viral supernatant via reverse transduction. Briefly, viral supernatant was added to a 6-well plate first. The cells were then counted and resuspended in D10 medium supplemented with 10 μg/mL of TransduceIT™ Transduction Reagent (Mirus Bio, Cat# MIR6620) and transferred to the wells pre-added with viral supernatant. The cells were incubated for 72 hours before flow cytometry sorting and/or antibiotic selection.

### Mammalian cell RNP nucleofection and plasmid transfection

Ribonucleoprotein (RNP) complexes were delivered into HEK293T cells via a 4D-Nucleofector® X Unit (Lonza, Cat# AAF-1003X) in a strip format using SF Cell Line 4D-Nucleofector™ X Kit S (Lonza, Cat# V4XC-2032). Purified PEs and CODEs were complexed with either pegRNA or cpegRNA to form RNP at 50 pmol protein: 200 pmol peg/cpegRNA ratio for 15 minutes at room temperature. Around 2×10^5^ cells were resuspended in Lonza SF buffer, mixed with the RNP, and electroporated using the program ED-130. The mixture was then incubated at 37°C and 5% CO^2^ for 10 minutes before adding to a 48-well plate pre-added with DMEM medium supplemented with 10% Fetal Bovine Serum (no antibiotic). Cells were harvested after 72 hours.

Plasmids encoding prime editors and chimeric editors and synthetic pegRNA/cpegRNA were delivered to HEK293T cells utilizing *Trans*IT-X2® Dynamic Delivery System (Mirus Bio, Cat# MIR6000). 24 hours prior to transfection cells were seeded at of 2×10^5^ cells per well in 24-well plates. Immediately prior to transfection, the media was replaced with antibiotic-free D10 media. 1500 ng of chimeric/prime editor plasmid, 750 ng of synthetic cpegRNA/pegRNA, and 500 ng of ngRNA plasmid (if applicable) were complexed for 20 minutes with 6 μL of *Trans*IT-X2 in 50 μL of Opti-MEM™ Reduced Serum Medium (Gibco, Cat# 31985062). After complexing, reactions were added dropwise to each well and harvested after 72 hours.

### Genomic DNA library preparation and targeted amplicon deep sequencing

Genomic DNA was extracted from the HEK293T and U2OS cells using the QuickExtract™ DNA Extraction Solution (Biosearch Technologies) system according to the manufacturer’s instructions. The DNA was then amplified in the first round of PCR using Q5 DNA Polymerase (NEB), and Illumina barcodes were appended during a second PCR. The products were then gel extracted, pooled together, and loaded on an Illumina MiSeqDx using a MiSeq Reagent Nano Kit v2 (Illumina, Cat# MS-101-1001) according to the manufacturer’s protocol. CRISPResso2 was used to determine the percentage of precise editing and indels^35^. Quantification window was defined by the parameter “-qwc” spanning 10 base pairs upstream and downstream flanking the targeting sequence in cases where there was no nicking guide and 10 base pairs flanking the targeting sequence and the nicking guide in the case where a nicking guide was used.

## Supporting information

List of Sequences

## Acknowledgments

We would like to thank members of Jain, Myhrvold, Toettcher, Adamson, and Burdine labs for insightful discussion and valuable suggestions. We are grateful for the Department of Molecular Biology, Omenn-Darling Bioengineering Institute, the Genomic Core at Princeton University, the NextGen DNA Sequencing core facility at the University of Florida (UF) Interdisciplinary Center for Biotechnology Research (ICBR), and UF Health Care Center at the University of Florida for their support. This work was financially supported in part by funds from the University of Florida (N.R.), the UF Herbert Wertheim College of Engineering (P.K.J.), Dinesh O. Shah endowed professorship (P.K.J.), the NIH-NIGMS Maximizing Investigator’s Research Award (MIRA) R35GM147788 (P.K.J.), National Institutes of Health (NIH) award number RM1HG009490 (B.A.), the Searle Scholars Program (B.A.), CHDI Foundation (B.A.), the China Scholarship Council (CSC) based on the April 2015 Memorandum of Understanding between the CSC and Princeton University (J.Y.), National Institutes of Health (NIH) award number U01DK127429 (J.E.T.), the New Jersey Commission on Cancer Research (NJCCR) #COCR24PRG007 (R.D.B.), National Institutes of Health (NIH) award number T32GM148739 (C.B.Y.), the Princeton Omenn-Darling Bioengineering Institute - Innovators (PBI2) program (L.N.), the Centers for Disease Control and Prevention award 75D30122C15113 (C.M.), and Princeton University (B.A., R.D.B., C.B.Y., C.M., L.N., J.Y., Y.J., and J.E.T.). The content is solely the responsibility of the authors and does not necessarily represent the official views of the National Institutes of Health. The findings and conclusions in this report are those of the authors and do not necessarily represent the official position of the CDC.

## Author Contributions

L.N., N.R., C.M., and P.K.J. conceptualized the ideas. L.N., N.R., C.M., and P.K.J. designed research. L.N., N.R., C.Y., B.P., Y.H., K.T.B., A.F., S.G.A., K.V.A., and C.M.D.P. carried out experiments. C.M., P.K.J., J.E.T., B.A., R.D.B., J.Y., and Y.J. helped troubleshoot the experiments and provide suggestions. L.N., N.R., B.P., and C.Y wrote the manuscript that was proofread and edited by all co-authors. The manuscript was approved by all authors.

## Competing Interests

L.N., N.R., B.P., C.M., and P.K.J. are listed as inventors on the patent applications related to the content of this work. P.K.J. is a co-founder of CasNx, LLC, Par Biosciences, LLC, and CRISPR, LLC. C.M. is a co-founder of Carver Biosciences. B.A. is an advisory board member with options for Arbor Biotechnologies and Tessera Therapeutics. B.A. holds equity in Celsius Therapeutics. J.Y. and B.A. have filed patent application(s) related to prime editing and/or other CRISPR-based technologies. J.E.T. is a scientific advisor for Prolific Machines and Nereid Therapeutics.

## Notes

### Summary of Updates

Figure 3 caption was missing sentences describing panels f and g. Figure 4 panel labeling was fixed.

