## Supplementary Information for "Efficient Genome Editing with Chimeric Oligonucleotide-Directed Editing"

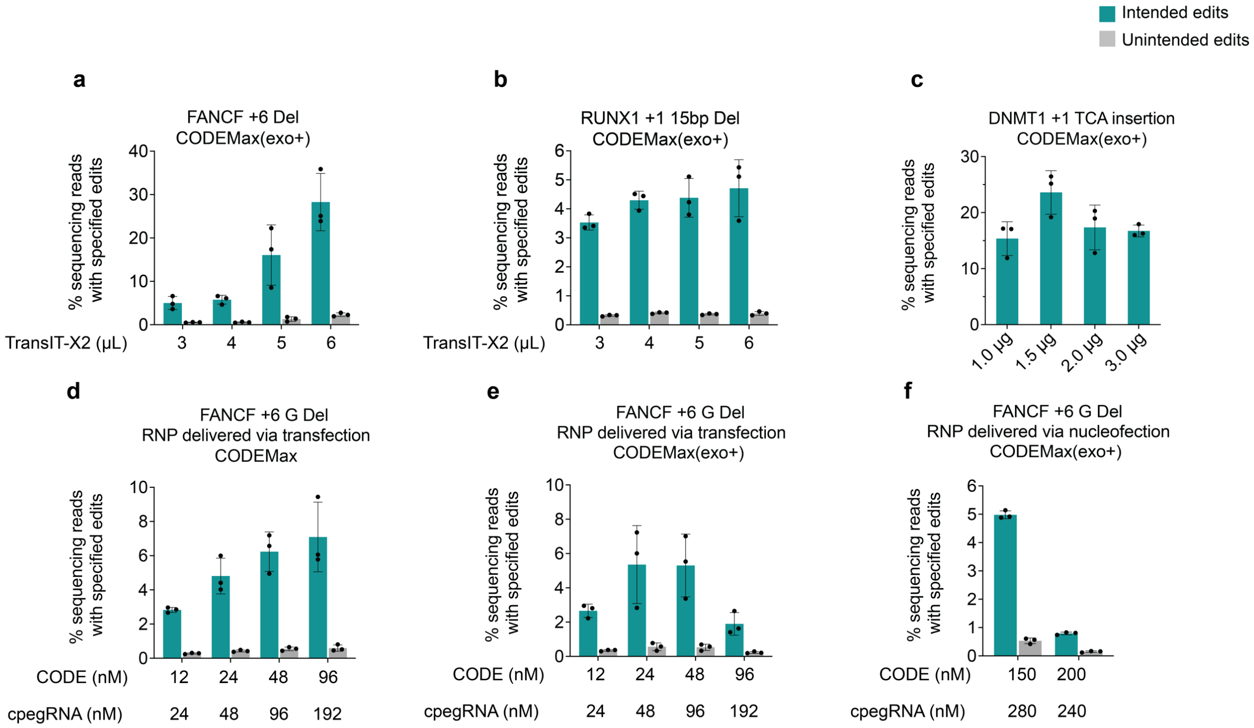


**Figure S1. General optimization of methods for chimeric oligonucleotide-directed editors.** (**a**) and (**b**) Effect of the volume of TransIT-X2 used during co-transfection of CODE encoding plasmid and cpegRNA at two different loci. (**c**) Effect of CODE encoding plasmid amount on editing efficiency. (**d**) and (**e**) Effect of the total RNP complex on editing efficiency via transfection. The ratio of cpegRNA to CODE was constant at 2:1. (**e**) Effect of the total RNP complex on editing efficiency via nucleofection. Error bars represent ± SD, where n = 3 technical replicates.

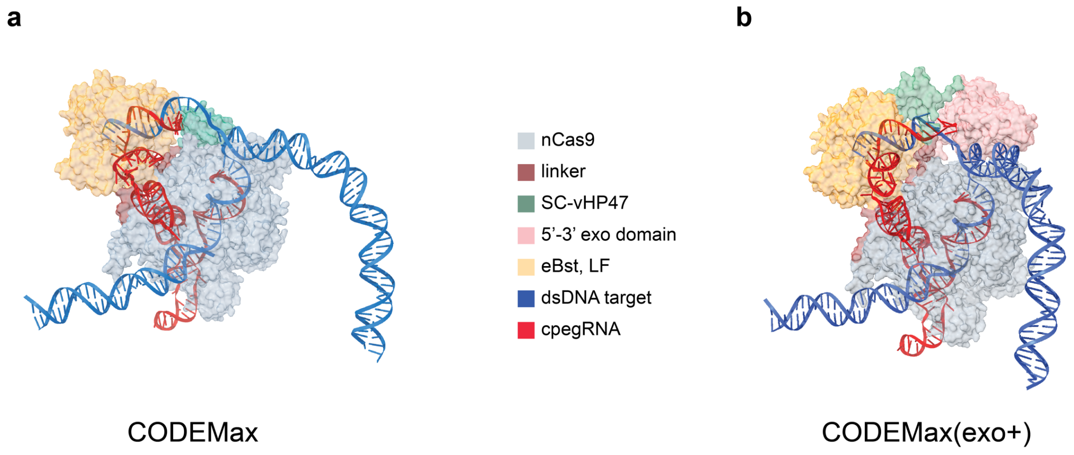


**Figure S2. Architecture of chimeric oligonucleotide-directed editors with domain organization.** AlphaFold 3 predicted structure of (**a**) CODEMax and (**b**) CODEMax(exo+) in complex with cpegRNA and target dsDNA[1]. cpegRNA modeled incorporated a C to T substitution at position +1 downstream of the nick site at the *HEK3* locus.

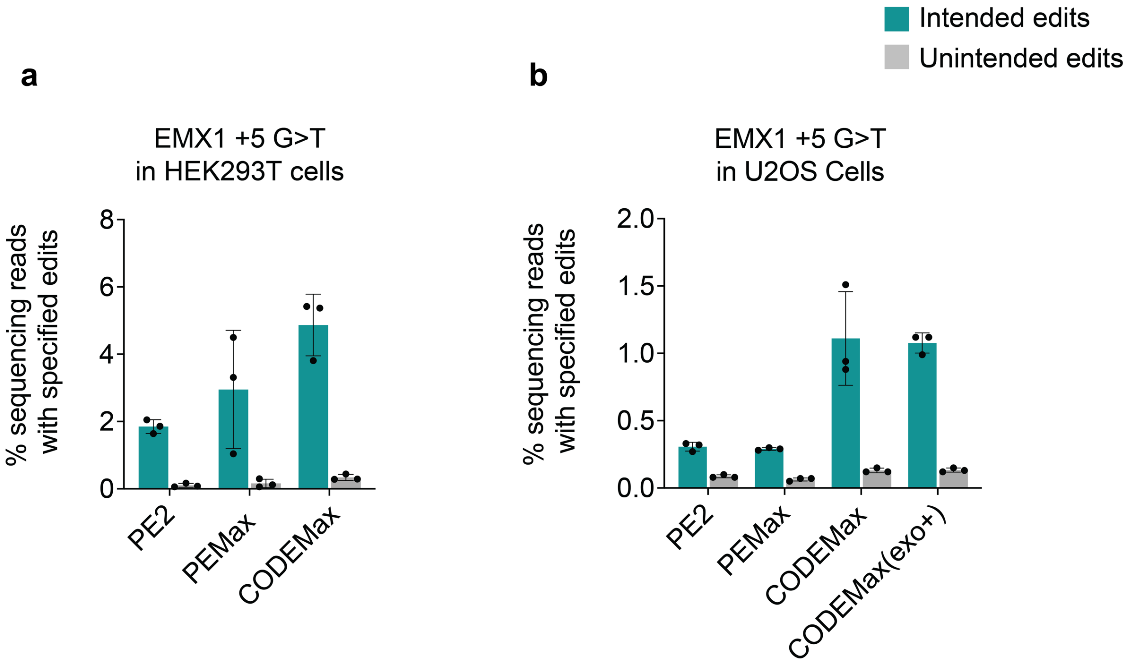


**Figure S3. Efficiency of prime editors and chimeric oligonucleotide-directed editors in different cell types.** (**a**) and (**b**) Efficiency of +5 G>T at the *EMX1* locus without ngRNA in HEK293T cells and U2OS cells, respectively. Error bars represent ± SD, where n = 3 technical replicates.

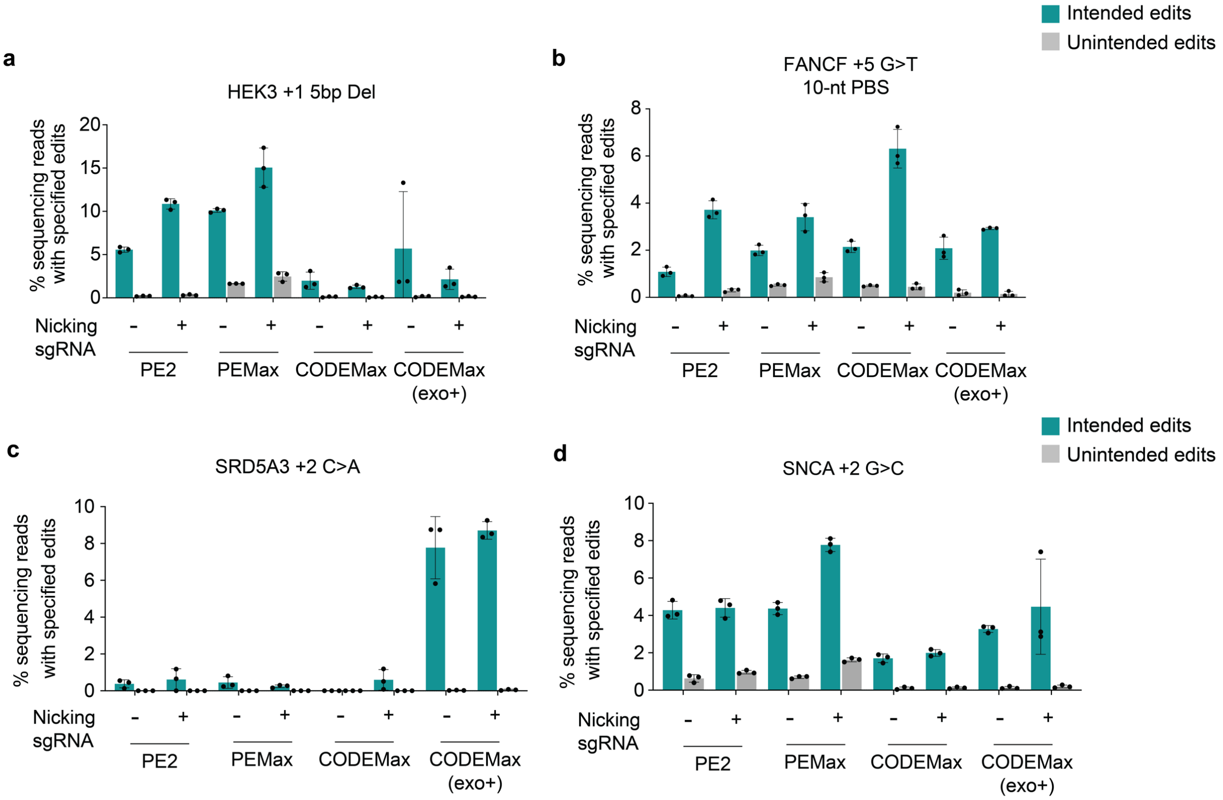


**Figure S4. Additional target sites for head-to-head comparison between prime editors and chimeric oligonucleotide-directed editors.** (**a**)-(**d**) Efficiency of intended and unintended modifications at *HEK3*, *FANCF*, and *SRD5A3*, and *SNCA* respectively. The data is complementary to Figure 4 in the main text. Error bars represent ± SD, where n = 3 technical replicates.

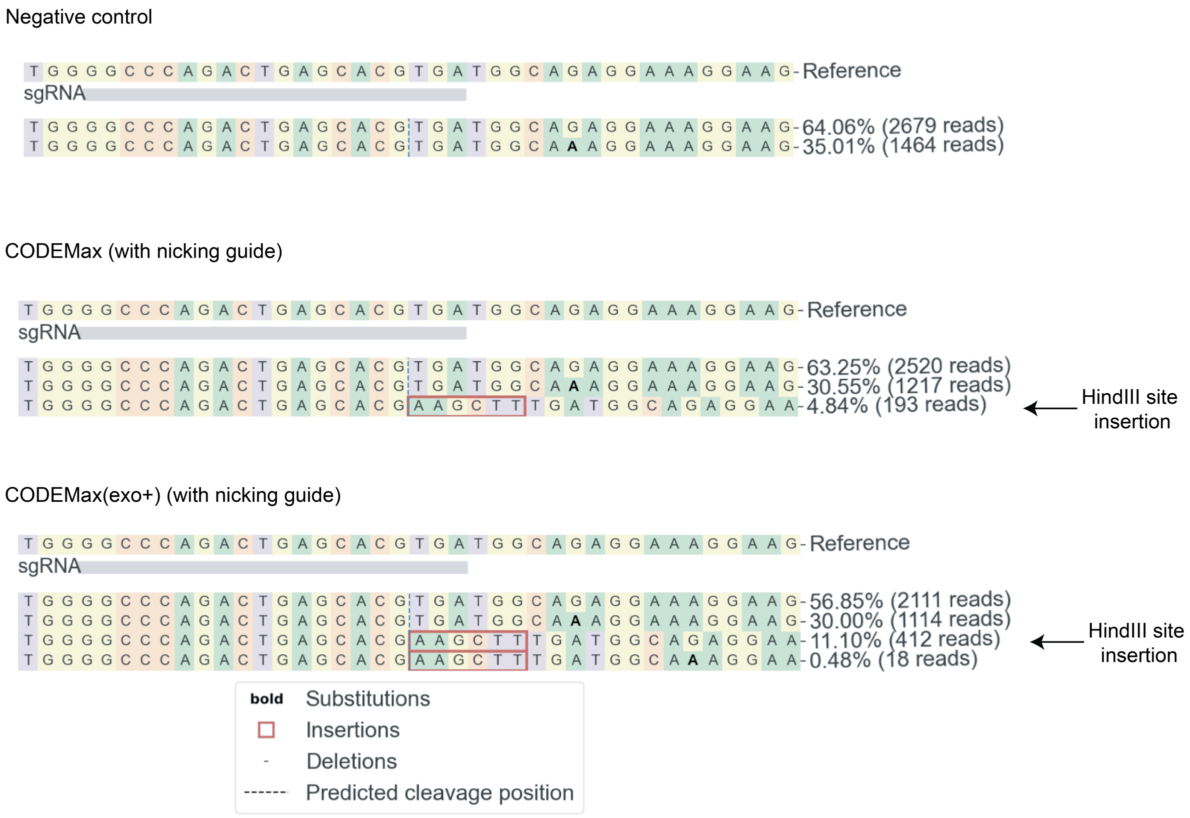


**Figure S5. Representative allele plot for insertion of HindIII at *HEK3* locus.** Allele plots with efficiencies and read counts were output by CRISPResso2[2].

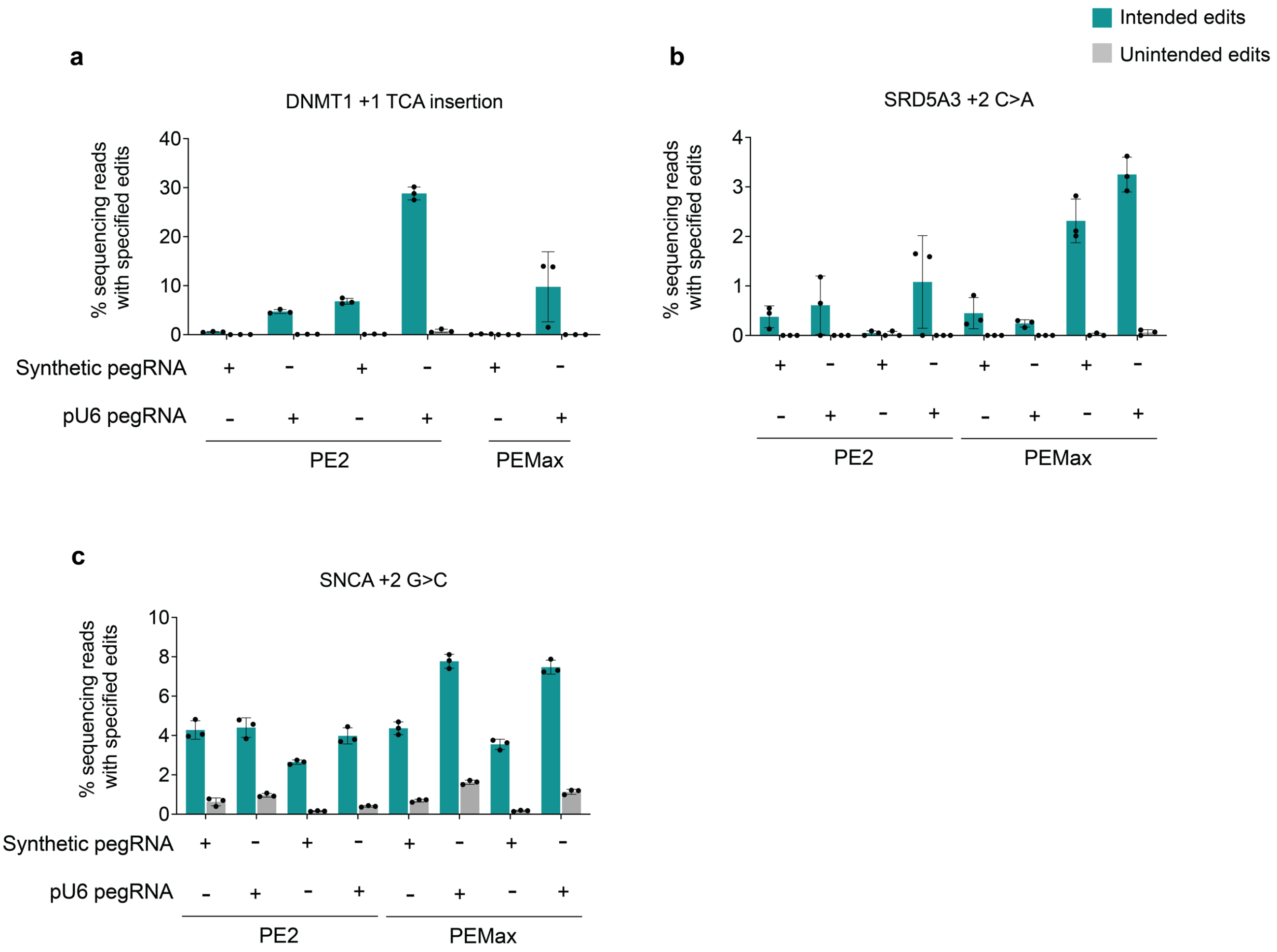


**Figure S6. Comparison of synthetic and pU6-promoter driven pegRNA delivery.**
(**a**) and (**b**) Head-to-head comparison of synthetic pegRNA and pU6-promoter driven pegRNA expression for edits at two endogenous loci for +1 TCA insertion at DNMT1, +2 substitution at SRD5A3, and +2 G>T at SNCA, respectively.

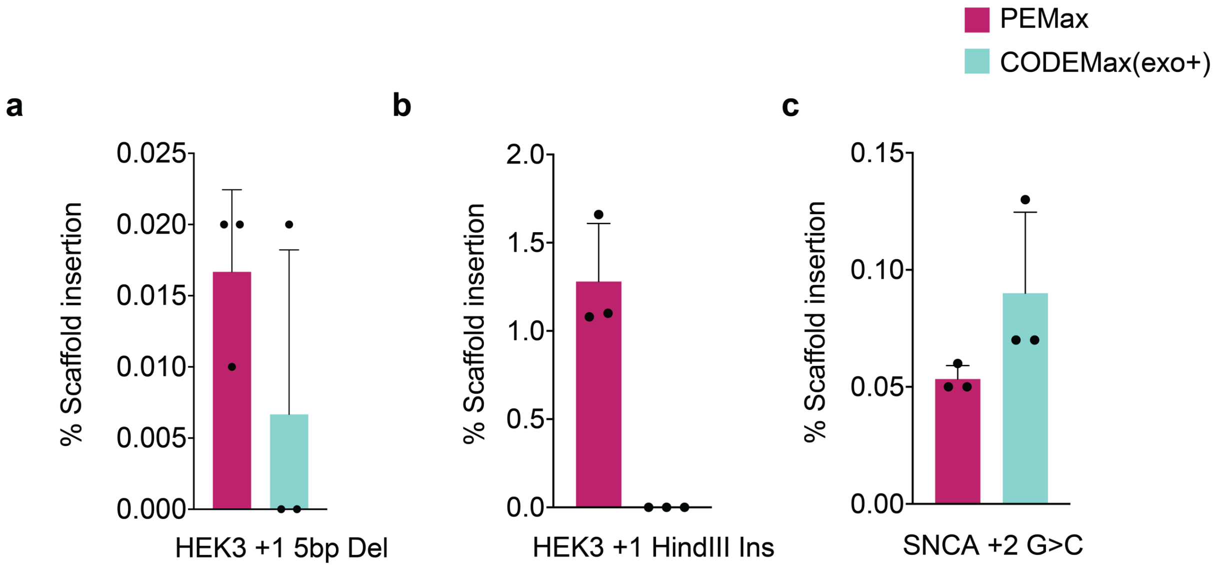


**Figure S7. Quantification of scaffold insertion for prime editors and chimeric oligonucleotide-directed editors for select edits.** (**a**), (**b**), and (**c**) Percent scaffold insertion quantified with CRISPResso2 at the *HEK3* locus for +1 5bp deletion and +1 HindIII insertion, and SNCA +2 G>C, respectively[2].

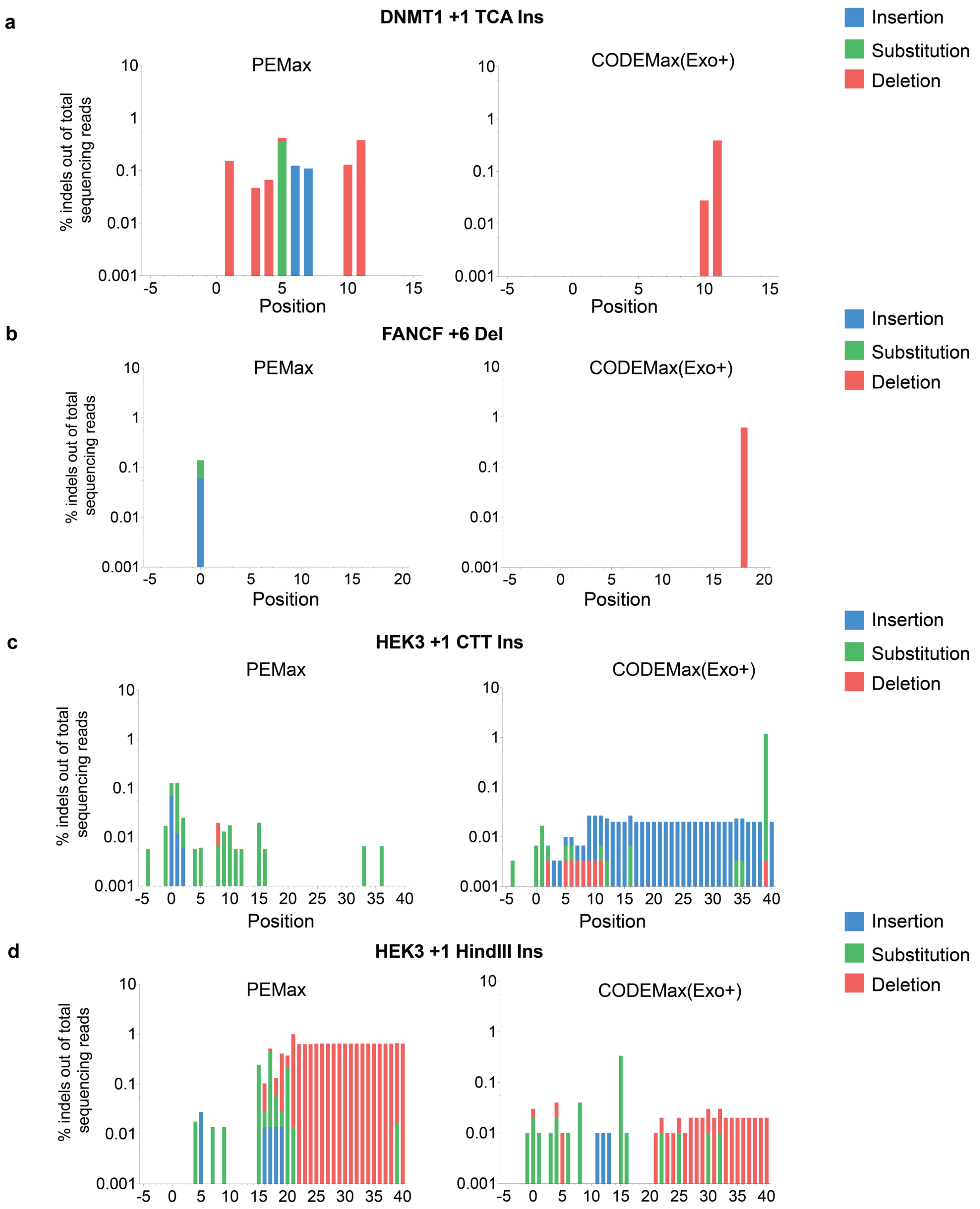


**Figure S8. Quantification of imprecise edits for prime editors and chimeric oligonucleotide-directed editors at select loci.** (**a**)-(**d**) Imprecise edit percentage and imprecise edit type as quantified by CRISPResso2 plotted against base position relative to nick site[2].
